# Augmentation of extracellular ATP synergizes with chemotherapy in triple negative breast cancer

**DOI:** 10.1101/2022.01.11.475873

**Authors:** Jasmine M Manouchehri, Jharna Datta, Natalie Willingham, Robert Wesolowski, Daniel Stover, Ramesh K Ganju, William Carson, Bhuvaneswari Ramaswamy, Mathew A Cherian

## Abstract

**Introduction:** Breast cancer affects two million women worldwide every year and is the most common cause of cancer-related death among women. The triple-negative breast cancer (TNBC) sub-type is associated with an especially poor prognosis because currently available therapies, fail to induce long-lasting responses. Therefore, there is an urgent need to develop novel therapies that result in durable responses. One universal characteristic of the tumor microenvironment is a markedly elevated concentration of extracellular adenosine triphosphate (eATP). Chemotherapy exposure results in further increases in eATP through its release into the extracellular space of cancer cells via P2RX channels. eATP levels are reduced by eATPases. Given that high concentrations of eATP are cytotoxic, we hypothesized that augmenting the release of eATP through P2RX channels and inhibiting extracellular ATPases would sensitize TNBC cells to chemotherapy.

**Methods:** TNBC cell lines MDA-MB 231, Hs 578t and MDA-MB 468 and non-tumorigenic immortalized mammary epithelial MCF-10A cells were treated with increasing concentrations the chemotherapeutic agent paclitaxel in the presence of eATPase inhibitors, specific agonists or antagonists of P2RXs with cell viability and eATP content being measured. Additionally, the mRNA, protein and cell surface expressions of the purinergic receptors P2RX4 and P2RX7 were evaluated in all examined cell lines via qRT-PCR, western blot, and flow cytometry analyses, respectively.

**Results:** In the present study, we observed dose-dependent declines in cell viability and increases in eATP in paclitaxel-treated TNBC cell lines in the presence of inhibitors of eATPases. These effects were reversed by specific antagonists of P2RXs. Similar results were observed with P2RX activators. All examined cell lines expressed both P2RX4 and P2RX7 at the mRNA, protein and cell surface levels.

**Conclusion:** These results reveal that eATP modulates the chemotherapeutic response in TNBC cell lines which could be exploited to enhance the efficacy of chemotherapy regimens for TNBC.

## Introduction

Breast cancer affects millions of women every year. At 47.8 new cases and 13.6 deaths per 100000 per year, it has the highest global incidence rate and is the most common cause of cancer-related mortality among women in 2020 (WHO, 2020). Patients with triple-negative breast cancer (TNBC) have a markedly worse outcome in comparison to other breast cancer subtypes due to the aggressive and rapidly progressive nature of the disease and lack of specific targeted therapies (Dent et al., 2009; Fisher et al., 2012; Ovcaricek, Frkovic, Matos, Mozina, & Borstnar, 2011). Hence, there is a critical need for more effective therapeutic strategies.

One universal characteristic of cancer is a marked elevation in extracellular adenosine triphosphate (eATP) (Fan et al., 2006; Pellegatti et al., 2008; Yegutkin, 2008). Under physiological conditions, the concentration of eATP is extremely low, in the 0-10 nanomolar (nM) range as compared to intracellular levels of 3 to 10 millimolar (mM), a difference of more than 10^6^-fold (Bakker et al., 2007). However, eATP concentrations are markedly elevated in neoplastic and inflamed tissues, into the range of 100s of micromoles/liter.

Purinergic P2 receptors (P2Rs), integral plasma membrane receptors activated by ATP, are divided into the ionotropic P2 (P2RX) and metabotropic P2 (P2RY) sub-types (Antonioli, Pacher, Vizi, & Hasko, 2013). P2XRs, of which there are seven sub-types, are ATP-gated ion channels that are inducibly permeable to cations. With prolonged activation, P2RX7 channels become non-selectively permeable, resulting in the diffusion of high molecular weight molecules such as ATP, IL-1β and IL-18 release and large molecular weight dye uptake, as well as K^+^ efflux, Na^+^ and Ca^2+^ influx, membrane phosphatidylserine-flip, membrane blebbing and cell death (Brandao-Burch, Key, Patel, Arnett, & Orriss, 2012; Chekeni et al., 2010; Ferrari et al., 2006; Mackenzie, Young, Adinolfi, & Surprenant, 2005; Yaron et al., 2015). Most P2RXs are activated by ATP concentrations in the nanomolar to low micromolar range, but P2RX7 activation requires millimolar concentrations of eATP. (Adinolfi et al., 2012; Gilbert et al., 2019; Haag et al., 2007). However, because P2RXs can homo and heterotrimerize to form functional channels with intermediate properties ATP-dependent interactions between P2RX4 and the C-terminus of P2RX7 can potentiate P2RX7-dependent cell death (Perez-Flores et al., 2015).

Extracellular ATP release occurs through a variety of mechanisms, including tumor necrosis and apoptosis, exocytosis, active efflux via ATP-binding cassette subfamily C member 6 (ABCC6) and the ankylosis gene product ANK and diffusion via P2RX7 and Pannexin1 channels (Akopova et al., 2012; Brandao-Burch et al., 2012; Chekeni et al., 2010; Elliott et al., 2009; Fader, Aguilera, & Colombo, 2012; Gobeil, Boucher, Nadeau, & Poirier, 2001; Jansen et al., 2014; Pellegatti, Falzoni, Pinton, Rizzuto, & Di Virgilio, 2005). Multiple pathways for eATP disposal have been described. These pathways hydrolyze nucleotides and limit the availability of nucleotides to activate nucleotide-specific P2Rs while increasing the concentration of extracellular nucleosides such as adenosine (Zimmermann, Zebisch, & Strater, 2012). There are four major classes of ecto-nucleotidases, including ecto-nucleoside triphosphate diphosphohydrolases (E-NTPDase), 5’ nucleotidases, ecto-nucleotide pyrophosphatases/phosphodiesterases (E-NPPase), and tissue non-specific alkaline phosphatases (TNAP) (Zimmermann et al., 2012). E-NTPDases, which are nucleotide specific, are believed to be the major degradative enzymes for eATP. Extracellular 5’nucleotidase, which is classified as CD73 as per the cluster of differentiation system, catalyzes the conversion of AMP to adenosine and inorganic phosphate. Thus, eATP levels result from a balance between numerous synthetic and secretory pathways and degradative and endocytic pathways.

When cancer cells are exposed to cytotoxic chemotherapy, there is a release of ATP and K^+^ ions through P2RXs such as P2RX7 and P2RX4 into the extracellular space along with an influx of Ca^2+^ ions (Di Virgilio & Adinolfi, 2017; Haag et al., 2007; Martins et al., 2009; Xia, Yu, Tang, Li, & He, 2015). Exposure of various epithelial cancer cell lines to elevated eATP in culture and xenografts results in growth arrest or cell death (Lertsuwan et al., 2017; Rapaport, 1988; Yoshihara et al., 2013). Notably, P2XR7 activation is a prerequisite for inflammasome activation, IL-1 and IL18 secretion, and a highly inflammatory form of programmed cell death known as pyroptosis, which can lead to bystander cell death and immune activation (Perez-Flores et al., 2015). In addition, ATP has been administered to patients with advanced cancers with minimal side effects, and ATP administered in mice was associated with inhibitory effects on cancer cells (Fontaine, 1996; Haskell, 1996; Lertsuwan et al., 2017).

Overall, these data suggest that the extracellular purinergic signaling pathway may be a promising target for cancer therapeutics. We hypothesized that increased eATP would increase the response to chemotherapy in TNBCs through the activation of P2RX channels, leading to increases in non-selective membrane permeability, the release of eATP and increased cancer cell death.

## Materials and Methods

### Cell Culture

Breast cancer cell lines MDA-MB 231 (ATCC HTB-26, RRID:CVCL_0062), MDA-MB 468 (ATCC HTB-132, RRID:CVCL_0419), Hs 578t (ATCC HTB-126, RRID:CVCL_0332), and HEK-293T ATCC Cat# CRL-3216, RRID:CVCL_0063)vwere maintained in DMEM (Corning) and supplemented with 10 % FBS (Cytiva), 1% MEM non-essential amino acids (Gibco), 1 mM sodium pyruvate (Gibco), 4 mM L-glutamine (Gibco) and antimicrobial agents (100 units/ml Penicillin, 100 μg/ml streptomycin, and 0.25 μg/ml amphotericin B) (Gibco).

Non-tumorigenic immortalized mammary epithelial MCF-10A cells (ATCC Cat# CRL-10317, RRID:CVCL_0598) were maintained in DMEM/F12 (Gibco) supplemented with 5% horse serum (Gibco), hydrocortisone (Sigma), epidermal growth factor (Sigma), cholera toxin (Sigma), insulin (Sigma) and antimicrobial agents. All cell lines were authenticated and were maintained in a humidified incubator at 37 °C and 5 % CO2.

### Drugs and chemicals

The following drugs and chemicals were used: dimethyl sulfoxide/DMSO (Sigma), ATP (Sigma), UTP (Sigma), paclitaxel (Calbiochem), sodium polyoxotungstate or POM-1 (Tocris), PSB 069 (Tocris), levamisole hydrochloride (Abcam), A438079 (Tocris), 5-BDBD (Tocris), ENPP1 inhibitor C (Cayman Chemical, Ann Arbor, MI), SBI-425 (MedChemExpress), etidronate disodium and ivermectin (Sigma). ATP and POM-1 were dissolved in nuclease-free water (Invitrogen); paclitaxel, A438079, Iso-PPADS, 5-BDBD, SBI-425, ENPP1 inhibitor C, levamisole hydrochloride, etidronate disodium (Sigma), and ivermectin were dissolved in DMSO.

### Cell viability and eATP assays

TNBC cell lines, MDA-MB 231, Hs 578t, MDA-MB 468 cells and non-tumorigenic immortalized mammary epithelial MCF-10A cells were plated on 96 well plates (Costar) at a high density of 25,000 cells/well and after 24 hours treated with paclitaxel, inhibitors, or ATP for 6 or 48 hours. PrestoBlue™ HS cell viability reagent (Invitrogen) was added to each well according to the manufacturer’s instructions. Fluorescence readings (excitation and emission ranges: 540–570 nm and 580–610 nm) were obtained using a Bioteck Synergy HT plate reader. ATP was measured in supernatants according to the protocol described by the ATPlite 1 step Luminescence Assay System (PerkinElmer). Luminescence readings were obtained from a Bioteck Synergy HT plate reader. The student’s t-test was applied to the applicable assays to ascertain significance.

### RNA analysis of P2RX4 and P2RX7

MDA-MB 231, Hs 578t, MDA-MB 468, cell lines and MCF-10A cells were maintained under standard conditions in subconfluent cultures. RNA was extracted via the TRIzol method (Invitrogen), and qRT-PCR was performed on a Bio-Rad T100 thermal cycler using the following exon-exon junction-spanning primers: for P2RX4, the forward primer TGGCGGATTATGTGATACCAGC and the reverse primer GTCGCATCTGGAATCTCGGG; for P2RX7, the forward primer GTGTCCCGAGTATCCCACC and the reverse primer GGCACTGTTCAAGAGAGCAG; and for GAPDH forward primer GTCGTATTGGGCGCCTGGTC and the reverse primer TTTGGAGGGATCTCGCTCCT. The student’s t-test was applied to the applicable assays to ascertain significance.

### Western blot analysis of P2RX4 and P2RX7

Total cell lysates were prepared in lysis buffer (50 mM Tris HCl at pH 8.0, 1.0 mM EDTA, 1% SDS, and 1% Igepal CA630) with a protease inhibitor cocktail (Thermo Scientific). The lysates were sonicated, placed on ice for 30 minutes, and spun at 10,000 rpm for 10 minutes at 4°C to collect the cleared supernatants for analysis. Protein quantification was performed using the Pierce BCA Protein Assay (Thermo Scientific) and absorbance readings taken at 595 nm. Protein samples were denatured with 4X Laemmli sample buffer (250 mM Tris-HCl, 8% SDS, 40% glycerol, 8% BME, and 0.06% Bromophenol Blue) at 98°C for 5 minutes and separated on 12% sodium dodecyl sulfate (SDS)-polyacrylamide gels (Invitrogen). Proteins were transferred to nitrocellulose membranes (Millipore) employing the wet transfer method (Bio-Rad, Hercules, CA). The membranes were blocked with non-fat milk at room temperature for an hour and incubated overnight at 4°C with a primary antibody: anti-P2RX4 (Cell Signaling Technology, Cat# 70659, RRID:AB_2799789) and anti-P2RX7 (Cell Signaling Technology, Cat# 13809, RRID:AB_2798319), diluted in 5% BSA (GoldBio) or 5% non-fat milk. The membranes were washed in TBS (0.15 M NaCl, 0.02 M Tris HCl, pH 7.4), incubated with horseradish peroxidase (HRP)-conjugated secondary anti-rabbit/mouse antibodies diluted in 5% non-fat milk (1:5000) for one hour and washed in TBS. The blots were analyzed using enhanced chemiluminescence Immobilon Western Chemiluminescent HRP Substrate (Millipore). The membranes were then stripped and re-probed for GAPDH (Cell Signaling Technology) as the internal loading control. Densitometry was performed on Image Studio (Licor). The student’s t-test was applied to the applicable assays to ascertain significance.

### Flow cytometry analysis of P2RX4 and P2RX7

MDA-MB 231, MDA-MB 468, Hs 578t, MCF-10A cells, and HEK 293T cells were maintained as previously described. HEK 293T were transfected with either P2RX4 or P2RX7 expression plasmids derived from pcDNA3.1 (RRID: Addgene_79663) using Lipofectamine 3000 (Thermo Fisher Scientific). Cells were detached with accutase (Thermo Fisher Scientific). One million cells were washed in PBS with 0.05% BSA, stained with P2RX7 –FITC (Sigma Aldrich, # P8997, RRID:AB_477416) antibodies or stained with rabbit IgG Isotype Control-FITC (Invitrogen, Cat# PA5-23092, RRID:AB_2540619), or with anti-P2RX4 (Cell Signaling Technology) plus goat anti-rabbit IgG (H+L) secondary antibody-FITC (Novus Biologicals, # NB 7168, RRID:AB_524413) or IgG isotype control plus secondary antibody-FITC in Flow Cytometry Staining Buffer (2% FBS, 0.02% sodium azide and PBS). Analysis was performed on BD FACS Fortessa and Flowjo software (RRID: SCR_008520). The student’s t-test was applied to the applicable assays to ascertain significance.

## Results

### Cell viability of ATP and UTP-treated cells and the impact of P2RX inhibitors on cell viability of ATP-treated cells

We examined the toxic effects of ATP using UTP as a control. We observed a dose-dependent loss of viability in TNBC MDA-MB 231, MDA-MB 468 and Hs 578t cell lines upon exposure to ATP but not UTP and not in non-tumorigenic immortalized mammary epithelial MCF-10A cells (**Figure 1**). For example, in ATP-treated (1 μmol/L) MDA-MB 231 cells, there was a 50% mean loss of viability (p=0.003 ATP vs vehicle; p=0.003 ATP vs UTP). Similar effects were observed in ATP-treated (1 μmol/L) Hs 578t cells with a 40% mean loss in viability (p=0.004 ATP vs vehicle; p=0.004 ATP vs UTP). For ATP-treated (1 μmol/L) MDA-MB 468 cells, there was a decrease in cell viability as compared to UTP-treated cells (p=0.05) but not significantly different from vehicle-treated control. There were no significant changes in viability in UTP-treated cells.

**Figure 1:**
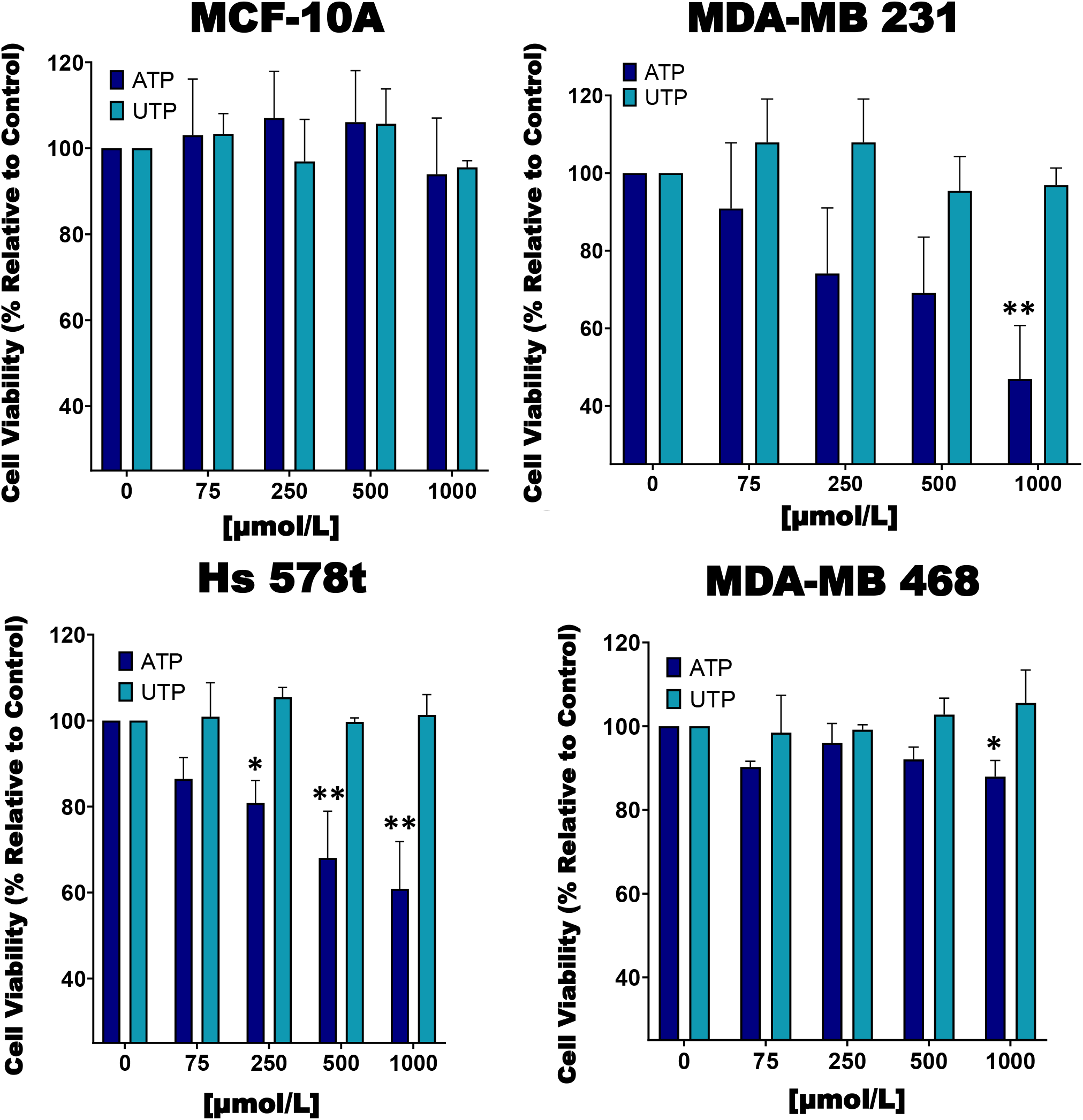
Cell viability of ATP and UTP-treated cells. TNBC MDA-MB 231, Hs 578t and MDA-MB 468 cell lines and non-tumorigenic immortalized mammary epithelial MCF-10A cells were treated for 48 hours with increasing concentrations of ATP or UTP, and cell viability was measured with the PrestoBlue HS assay. Error bars represent standard deviations calculated from three independent experiments performed in triplicate. Y axis begins at 25%. The student’s t-test was applied to the applicable assays to ascertain significance. * represents p<0.05 and ** represents p<0.01.

We next treated the cells with various purinergic receptor antagonists to determine whether P2RX receptors mediate the effects of eATP on cell viability (**Figure 2**). As with figure 1, we did not see any change in cell viability in ATP-treated MCF-10A cells and therefore, did not see any additional changes with exposure to P2RX antagonists. However, for ATP-treated TNBC cells, in the absence of inhibitors we saw decreases in viability that were attenuated by P2RX inhibitors, thus decreasing their sensitivity to inhibition by ATP. For example, in ATP-treated (1 μmol/L) MDA-MB 231 cells, when compared to vehicle addition, there was a 15% improvement in cell viability when exposed to the non-specific P2RX inhibitor Iso-PPADS (20 μmol/L) (p=0.04), a 30 % improvement in cell viability when exposed to the P2RX7 inhibitor A438079 (20 μmol/L) (p=0.01), and a 30% improvement in cell viability when exposed to the P2RX4 inhibitor 5-BDBD (20 μmol/L) (p=0.01). Similar effects were seen in ATP-treated (1 μmol/L) Hs 578t cells: when compared to vehicle addition, there was a 33% improvement in cell viability when exposed to Iso-PPADS (p=0.01), 30 % improvement in cell viability when exposed to A438079 (p=0.01), and a 32% improvement in cell viability when exposed to 5-BDBD (p=0.01). For ATP-treated (1 μmol/L) MDA-MB 468 cells as compared to vehicle-addition, there was a 23% improvement in cell viability when exposed to Iso-PPADS (p=0.02), a 10% improvement in cell viability when exposed to A438079 (p=0.04), but no significant improvement in cell viability when exposed to 5-BDBD.

**Figure 2:**
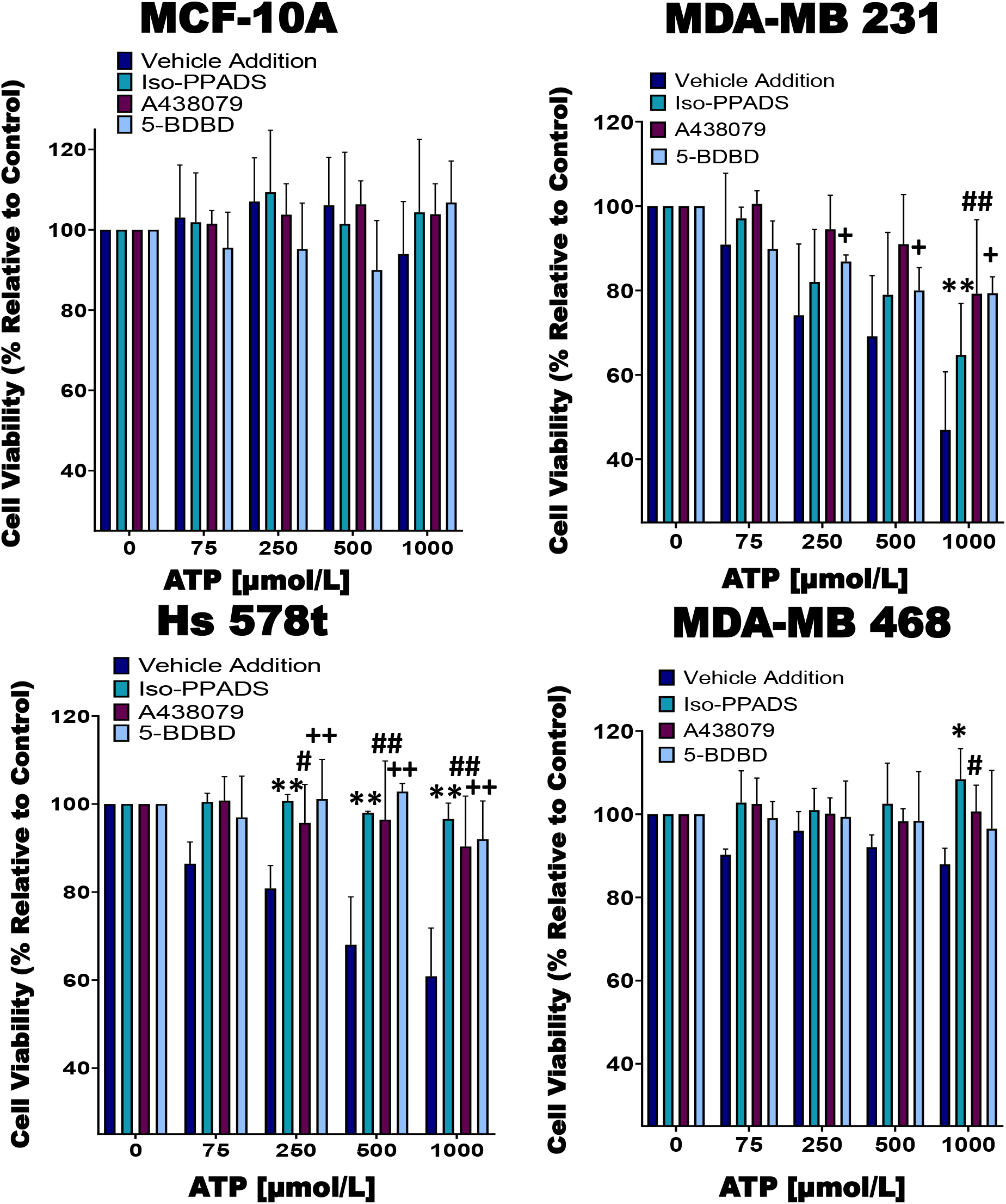
Cell viability of ATP-treated cells in the presence of P2RX inhibitors. TNBC cell lines and MCF-10A cells were treated for 48 hours with increasing concentrations of ATP in the presence of the P2RX inhibitor Iso-PPADS (20 μmol/L), the P2RX7 inhibitor A438079 (20 μmol/L) or the P2RX4 inhibitor 5-BDBD (20 μmol/L) or vehicle addition, and cell viability was measured using the PrestoBlue HS assay. Error bars represent standard deviations calculated from three independent experiments performed in triplicate. Y axis begins at 25%. The student’s t-test was applied to the applicable assays to ascertain significance.* represents p<0.05 and ** represents p<0.01 for Iso-PPADS; # represents p<0.05 and ## represents p<0.01 for A438079; + represents p<0.05 and ++ represents p<0.01.

### Examining the effects of eATPase inhibitors on cell viability and eATP release

We next studied the effects of combinations of eATPase inhibitors with chemotherapy (paclitaxel) to determine their effects on the efficacy of chemotherapy. For these experiments, all the cell lines were treated for six hours to simulate the duration of systemic exposure in patients (**Figure 3**). For this reason, we did not see changes in the viability of cells treated with paclitaxel alone. Also, we did not see significant changes in cell viability in paclitaxel-treated MDA-MB 231 cells in the presence of the E-NTPDase inhibitor POM-1 when compared to vehicle addition with paclitaxel. However, there was a 30% mean decrease in cell viability in the presence of the E-NTPDase inhibitor PSB 069 (p=0.01) and a 34% mean decrease in cell viability in the presence of the ENPPase inhibitor ENPP1 inhibitor C (p=0.01) when compared to vehicle addition. Similarly for paclitaxel-treated Hs 578t cells in the presence of POM-1, there was a 35% loss of viability (p=0.01) in the presence of PSB 069, a 27% loss of viability (p=0.02) and in the presence of ENPP1 inhibitor C, a 25% loss of viability (p=0.02). For paclitaxel-treated (100 μmol/L) MDA-MB 468 cells there was a 50% decrease in cell viability in the presence of PSB 069 as compared to vehicle addition (p=0.01) but no significant loss of viability in the presence of POM-1 or ENPP1 C inhibitor. However, there was no significant change in the viability of treated MCF-10A cells (**Figure 3A**). We also confirmed that under these experimental conditions, treatment with the eATPase inhibitors alone did not significantly change cell viability in any of the cell lines (**Supplemental Figure 1**). Therefore, PSB 069 most potently and consistently decreased the viability of TNBC cell lines when combined with paclitaxel.

**Figure 3:**
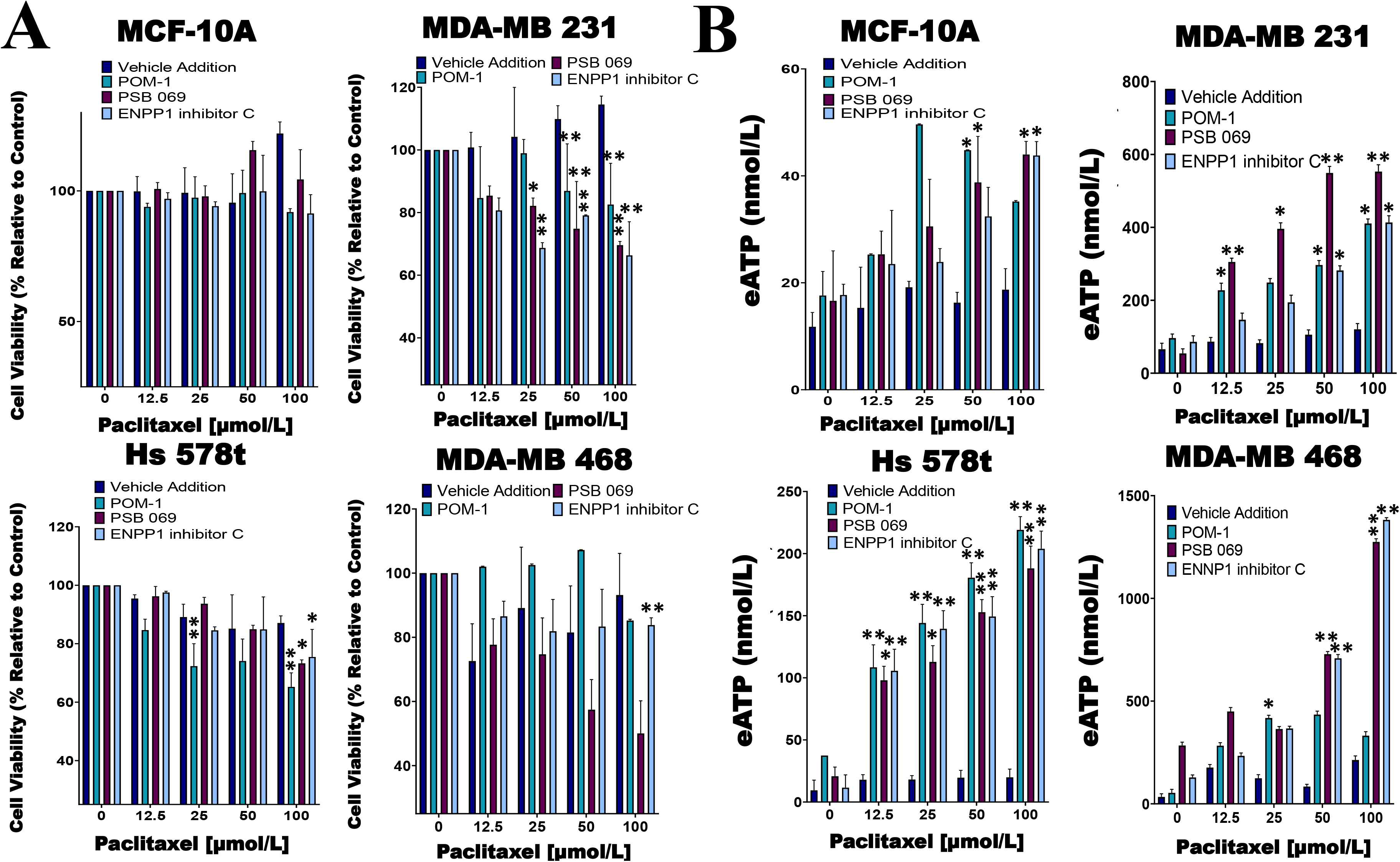
Comparing eATP release from paclitaxel-treated cells in the presence of inhibitors or vehicle addition. **(A)** TNBC and MCF-10A cell lines were treated with increasing concentrations of paclitaxel and the nucleoside phosphohydrolase inhibitors POM-1 (E-NTPDase inhibitor, 10 μmol/L), PSB 069 (E-NTPDase inhibitor, 10 μmol/L), ENNP1 inhibitor C (ENPP1 inhibitor, 10 μmol/L) or vehicle addition for six hours, and cell viability was measured using the PrestoBlue HS assay. Standard deviation was calculated from three independent experiments performed in triplicate. Y axis begins at 25%. **(B)** eATP concentrations were measured in the supernatants of TNBC and MCF-10A cell lines after six hours of treatment with increasing concentrations of paclitaxel and nucleoside phosphohydrolase inhibitors or vehicle addition. Standard deviation was calculated from three independent experiments performed in triplicate. The student’s t-test was applied to the applicable assays to ascertain significance.* represents p<0.05 and ** represents p<0.01.

In the same experiments, we measured the amount of eATP in the supernatants of chemotherapy-treated cells (**Figure 3B**). Treating cells with 100 μmol/L paclitaxel alone produced quite modest increases in eATP that were generally not statistically significant when compared to vehicle control: for MDA-MB 231, eATP increased to 120 nmol/L, for Hs 578t, eATP increased to 20 nmol/L, for MDA-MB 468, eATP increased to 213 nmol/L, and for MCF-10A, eATP increased to 18 nmol/L. However, in the presence of inhibitors, we saw significant increases in eATP levels. For instance, in paclitaxel-treated (100 μM) MDA-MB 231 cells, the concentration of eATP increased upon treatment with POM-1 (450 nmol/L; p=0.02 vs vehicle addition), with PSB 069 (550 nmol/L; p=0.001 vs vehicle addition); and with ENPP1 inhibitor C (410 nmol/L; p=0.02 vs vehicle addition). Similarly, for paclitaxel-treated (100 μmol/L) Hs 578t cells with POM-1 (219 nmol/L; p=0.01 vs vehicle addition), with PSB 069 (188 nmol/L; p=0.02 vs vehicle addition and with ENPP1 inhibitor C (204 nmol/L; p=0.01 vs vehicle addition). For paclitaxel-treated (100 μmol/L) MDA-MB 468 cells, the concentration of eATP increased with PSB 069 (1.3 μM /L; p=0.003 vs vehicle addition); and with ENPP1 inhibitor C (1.4 μmol/L; p=0.003 vs vehicle addition), but not statistically significant increase with POM-1 (300 nmol/L). Thus, ENTPDase and NPPase inhibitors significantly increased eATP release upon chemotherapy treatment.

### Examining the effects of P2RX inhibitors on the E-NTPDase inhibitor-induced exaggerated loss of cell viability and eATP release

Of the eATPase inhibitors tested, we consistently saw an exaggerated loss of cell viability with the E-NTPDase inhibitor PSB 069. Therefore, we sought to determine if the exaggerated loss of cell viability in the presence of PSB 069 is dependent on eATP induced activation of P2RX4 or P2RX7 (**Figure 2**). We did see reversal of the effects of PSB 069 on cell viability and eATP release upon exposure to both the P2RX7 inhibitor A438079 and the P2RX4 inhibitor 5-BDBD (**Figure 4A**). In paclitaxel (100 μmol/L) treated MDA-MB 468 cells, there was a 34% improvement in viability for the combination of A438079 with PSB 069 when compared to vehicle addition with PSB 069 (p=0.01) and a 27% improvement in cell viability in the presence of 5-BDBD and PSB 069 when compared to vehicle addition with PSB 069 (p=0.03).

**Figure 4:**
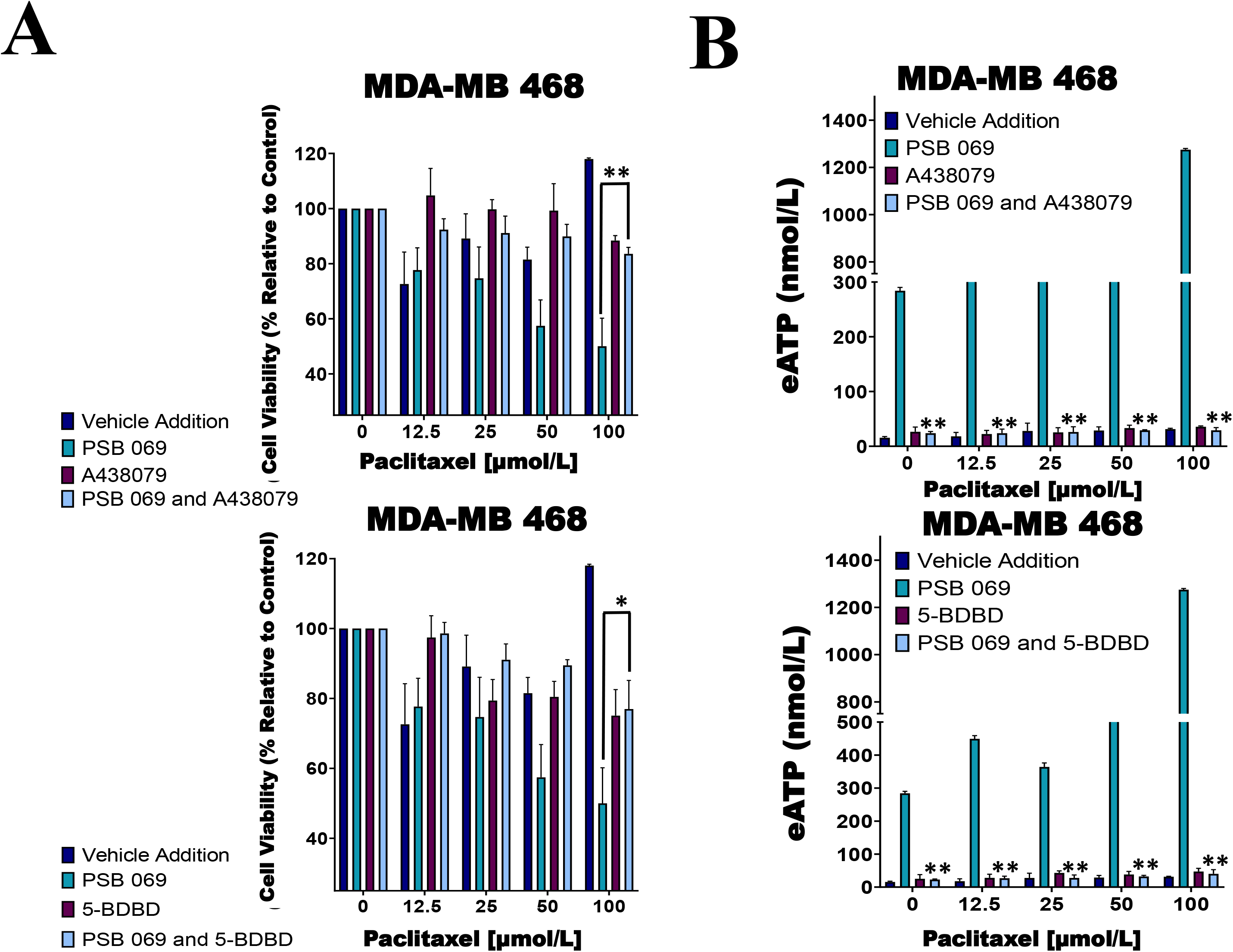
Examining the influence of P2RX inhibitors in combination with E-NTPDase inhibitor on cell viability and eATP release in paclitaxel-treated cells. (A) Paclitaxel-treated breast cancer MDA-MB 468 cell lines were treated for six hours with P2RX7 inhibitor A438079 (20 μmol/L) or P2RX4 inhibitor 5-BDBD (20 μmol/L) in the presence or absence of PSB 069 (10 μmol/L), and cell viability was measured by applying PrestoBlue HS assay. Standard deviation was calculated from three independent experiments performed in triplicate. We used that same values for both graphs for vehicle addition and PSB 069. Y axis begins at 25%. **(B)** eATP concentrations were measured in the supernatants of paclitaxel-treated MDA-MB 468 cells after six hours of treatment. Standard deviation was calculated from three independent experiments performed in triplicate. We used that same values for both graphs for vehicle addition and PSB 069. The student’s t-test was applied to the applicable assays to ascertain significance.* represents p<0.05 and ** represents p<0.01 for A438079 and PSB 069 or 5-BDBD and PSB 069 when compared to PSB 069.

In the same experiments, we determined the effects of A438079 and 5-BDBD on PSB 069- and paclitaxel (100 μmol/L)-induced eATP release in MDA-MB 468 cells **(Figure 4B)**. There was a decrease in eATP from 1.3 μmol/L to 30 nmol/L when A438079 was combined with PSB 069 (p=0.001 when comparing A438079 with PSB 069 to vehicle addition with PSB 069). There was a decrease in eATP from 1.3 μmol/L to 40 nmol/L when 5-BDBD was combined with PSB 069 (p=0.001 when comparing 5-BDBD with PSB 069 to vehicle addition with PSB 069). These results show that the exaggerated loss of cell viability observed when PSB 069 is combined with paclitaxel is dependent on the activation of P2RX4 and P2RX7 by eATP.

### Analyzing the effect of phosphatase inhibitors on cell viability and ATP release from chemotherapy-treated cells-

Previous reports indicate that tissue non-specific alkaline phosphatase also metabolizes eATP. We sought to ascertain if two tissue non-specific alkaline phosphatase inhibitors (SBI 425 and levamisole hydrochloride) could augment the effects on cell viability and eATP release in paclitaxel-treated cells while using a protein tyrosine phosphatase inhibitor, etidronate disodium, as a control. Although we observed substantial changes in eATP upon treatment of the TNBC cell lines, there was no significant change in cell viability **(Supplemental Figure 2)**.

### Evaluating the impact of a P2RX activator on cell viability and ATP release in chemotherapy-treated cells

Previous research had shown that ivermectin is a P2RX4 and P2RX7activator (Khakh, Proctor, Dunwiddie, Labarca, & Lester, 1999; Nörenberg et al., 2012). Hence, we examined the effects of ivermectin on eATP and cell viability in chemotherapy-treated MCF10A cells and TNBC cell lines. We observed significant decreases in cell viability in paclitaxel and ivermectin-treated TNBC cell lines but not in MCF-10A cells (**Figure 5A**). As an example, for paclitaxel (100 μmol/L) treated cells in the presence of ivermectin (20 μmol/L) as compared to vehicle addition, MDA-MB 231, Hs578t and MDA-MB 468 cells showed 35 % (p=0.01), 38% (p=0.01) and 50% (p=0.001) mean decreases in cell viability, respectively.

**Figure 5:**
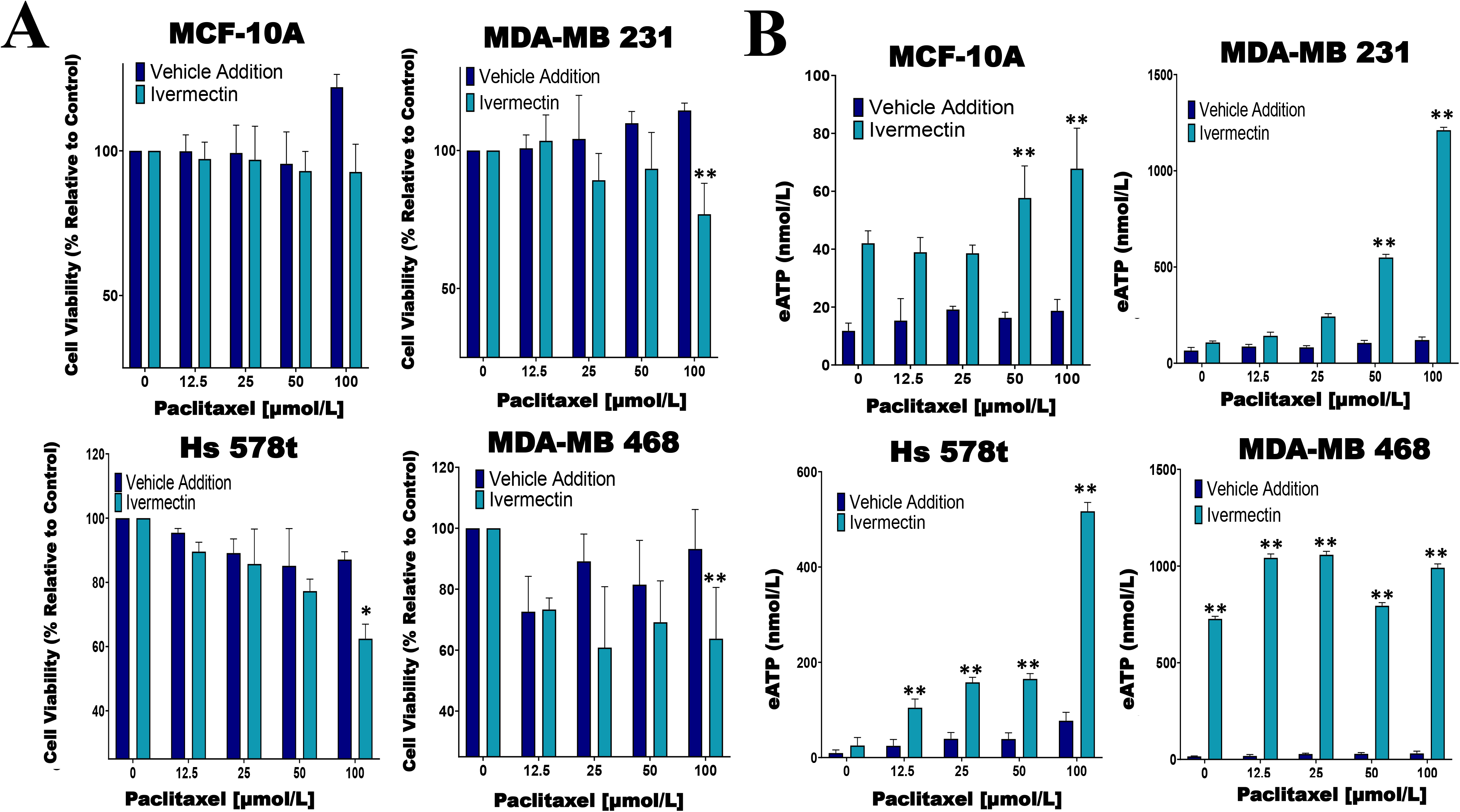
Determining relative eATP content and cell viability in paclitaxel-treated cells in the presence of ivermectin or vehicle addition. **(A)** The graphs represent cell viability as measured using the Presto Blue HS assay +/- standard deviation from three independent experiments performed in triplicate in TNBC and MCF-10A cell lines after six hours of treatment with increasing concentrations of paclitaxel and the P2RX4 and P2RX7 activator ivermectin (10 μmol/L) or vehicle addition. Y axis begins at 25%. **(B)** eATP content was measured in the supernatants of paclitaxel-treated TNBC and MCF-10A cell lines in the presence of the P2RX4 and P2RX7 activator ivermectin (10 μmol/L) or vehicle addition. The student’s t-test was applied to the applicable assays to ascertain significance.* represents p<0.05 and ** represents p<0.01.

In the same experiments, we also looked at eATP release upon exposure to the combined treatment of ivermectin and paclitaxel. For paclitaxel-treated (100 μmol/L) cells there were increases in eATP release in the presence of ivermectin when compared to the vehicle addition, however, these increases were much more dramatic in the TNBC cell lines (**Figure 5B**). As an example, for MDA-MB 231 cells in the presence of ivermectin, eATP increased from 120 nmol/L (vehicle addition) to 1.2 μmol/L (p=0.002), for Hs 578t cells eATP increased from 20 nmol/L (vehicle addition) to 517 nmol/L (p=0.001) and for MDA-MB 468 cells eATP increased from 213 nmol/L (vehicle addition) to 1 μmol/L (p=0.002). Therefore, ivermectin potentiated the effects of paclitaxel on TNBC cell lines.

### Expression of P2RX4 and P2RX7 in TNBC cell lines

We next sought to assess the expression of P2RX4 and P2RX7 mRNA and protein. qRT-PCR was performed on TNBC and MCF-10A cells with specific exon-exon junction-spanning primers for *P2RX4, P2RX7* and *GAPDH*, and fold change was calculated relative to the expression of the receptors in MCF-10A cells (**Figure 6A**). Some TNBC cell lines expressed more *P2RX4* mRNA in comparison to MCF-10A cells: MDA-MB 231 (5-fold; p=0.0012 and MDA-MB 468 (10-fold; p=0.0001); whereas Hs 578t cells expressed levels that were not significantly different (p>0.05). MDA-MB 231 and Hs 578t cells expressed significantly less *P2RX7* mRNA when compared to MCF-10A cells (p=0.0001 for both); whereas, MDA-MB 468 cells expressed 25-fold more *P2RX7* mRNA when compared to MCF-10A cells (p=0.0006).

**Figure 6:**
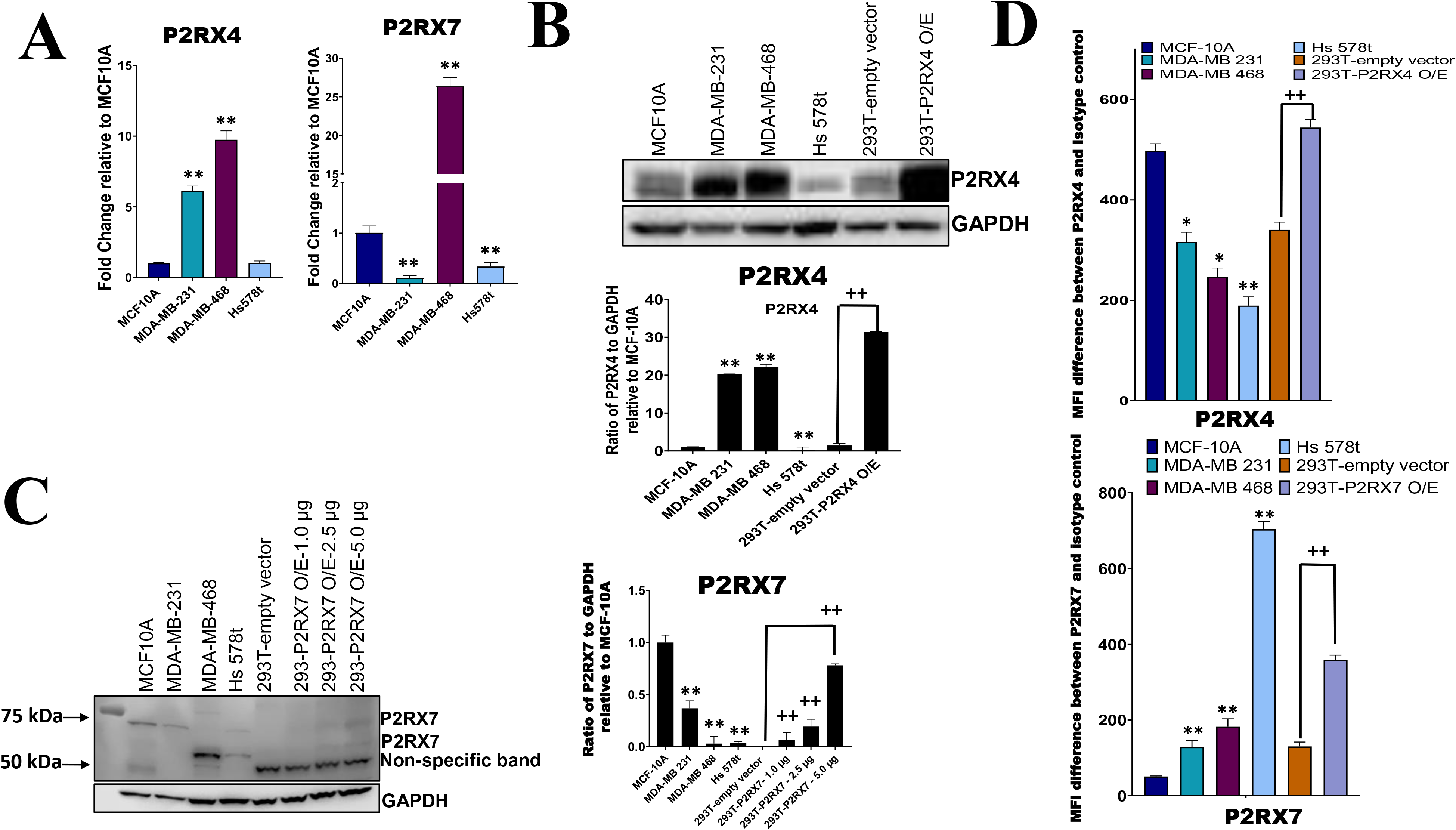
mRNA and protein expression analysis of P2RX4 and P2RX7 for all cell lines. **(A)** qRT-PCR was performed on mRNA of TNBC cell lines and MCF-10A cells using specific primers for P2RX4 and P2RX7. * represents p<0.05 and ** represents p<0.01. TNBC cell lines, MCF-10A cells, and HEK 293T cells transfected with P2RX4 or P2RX7 as positive controls were probed for **(B)** P2RX4 and **(C)** P2RX7, and GADPH was used as a loading control for western blot analysis repeated twice. HEK 293T cells transfected with P2RX7 were loaded at increasing protein concentrations of 1.0 μg, 2.5 μg, and 5.0 μg combined with lysates of control vector-transfected cells to keep the total loaded protein the same in each lane. Densitometry analysis was performed using Image Studio on the 75 kDa P2RX7 band. The student’s t-test was applied to the applicable assays to ascertain significance. * represents p<0.05 and ** represents p<0.01 relative to MCF-10A; + represents p<0.05 and ++ represents p<0.01 relative to HEK293-empty vector transfected. **(D)** The calculated difference in mean fluorescence intensity (MFI) values between TNBC cell lines, MCF-10A cells, and HEK 293T cells transfected with P2RX4 or P2RX7 as positive controls stained with P2RX4 or P2RX7 specific antibody and the isotype control for the different cell lines examined. * represents p<0.05 and ** represents p<0.01 relative to MFI difference in MCF-10A cells; + represents p<0.05 and ++ represents p<0.01 relative to MFI difference in HEK293-empty vector transfected. O/E represents overexpressed.

**Figure 7:** Schematic displaying our proposed model for ATP release. Our proposed model suggests that ivermectin activates P2RX4 and P2RX7 leading to the release of ATP and the more ATP that accumulates extracellular can promote cell death especially in the presence of paclitaxel. In addition, the breakdown of ATP can be prevented in the presence of E-NTPDase inhibitors POM-1 or PSB 069. However, the release of ATP can be prevented in the presence of P2RX4 inhibitors 5-BDBD or Iso-PPADS or P2RX7 inhibitors A438079 or Iso-PPADS.

Western blot analysis was performed on TNBC cell lines, MCF-10A cells and HEK 293T cells transfected with either a P2RX4 or P2RX7 expression plasmid as positive controls, probing for P2RX4 and P2RX7 with GAPDH as the internal loading control (**Figure 6B**). Two of three TNBC cell lines expressed more P2RX4 protein when compared to MCF-10A cells when assessed by semi-quantitative densitometry: MDA-MB 231 (20-fold; p=0.001), MDA-MB 468 (22-fold; p=0.001) while Hs 578t cells expressed significantly less P2RX4 protein (p=0.01). We separately probed for P2RX7 protein in the same cell lines. We did detect specific bands corresponding to the full-length glycosylated P2RX7A isoform (75kDa) in the MCF-10A cells but significantly less protein was detected in MDA-MB 231 and Hs 578t cells while the 69 kDa was detected in MDA-MB 468 and Hs 578t cells; we did detect specific bands at 69 and 75 kDa in transfected 293T cells that increased in intensity with increasing mass of transfected P2RX7 plasmid. This plasmid incorporates the cDNA for the full-length P2RX7A isoform. The P2RX7 protein includes 5 N-linked glycosylation sites. Thus, the 69 kDa band corresponds to the unglycosylated form of the protein and the 75 kDa band likely represents the fully glycosylated form of the protein.

Given that unglycosylated form may not represent the membrane-localized fraction of a protein, we used flow cytometry to quantitate the expression of P2RX7 and P2RX4 at the cell surface. Flow cytometry analysis was performed on TNBC, MCF-10A, and HEK 293T cells transfected with P2RX4 as a positive control probing for cell surface expression P2RX4 (**Figure 6D**) using antibodies that target the extracellular domains of P2RX4. MDA-MB 231, Hs 578t and MDA-MB 468 expressed significantly less cell surface P2RX4 in comparison to MCF-10A cells. The calculated mean fluorescence intensity (MFI) difference between the P2RX4 specific and isotype control antibody for MCF-10A cells was 498, for MDA-MB 231 cells 316 (p=0.05 when compared to MFI difference for MCF-10A), for MDA-MB 468 cells 246 (p=0.02), and Hs 578t cells 189 (p=0.01).

We performed flow cytometry analysis on TNBC, MCF-10A, and HEK 293T cells transfected with P2RX7 as a positive control probing for cell surface expression P2RX7 (**Figure 6D**) using antibodies that target the extracellular domains of P2RX7. MDA-MB 231, Hs 578t and MDA-MB 468 expressed significantly more cell surface P2RX7 in comparison to MCF-10A cells. The calculated mean fluorescence intensity (MFI) difference between P2RX7 and the isotype control for MCF-10A cells was 51, for MDA-MB 231 cells 129 (p=0.003 when compared to MFI difference to MCF-10A), for MDA-MB 468 cells 182 (p=0.002), and for Hs 578t cells 703 (p=0.0001). Thus, all cell lines expressed both receptors at the cell surface when measured by flow cytometry.

Additionally, all cell lines were treated with 100 μmol/L paclitaxel and the cell surface expressions of P2RX4 and P2RX7 were examined with the calculated difference in MFI values from P2RX4 or P2RX7 stained from the corresponding isotype control (**Figure S4**). Upon treatment, cell surface expression of P2RX7 increased significantly in some TNBC cell lines but this was not a consistent effect.

## Discussion

Chemotherapy by itself fails to ablate metastatic TNBC. This is still an incurable disease with the shortest median overall survival among breast cancer subtypes. Thus, new therapeutic strategies are urgently required. Extracellular ATP, in the high micromolar to millimolar range, induces cytotoxicity in cancer cell lines. Chemotherapy is known to induce increases in eATP. We hypothesized that interventions that augment chemotherapy-induced increases in eATP would increase cancer cell death. In our study, the commonly used chemotherapeutic agent paclitaxel was applied to TNBC cells and non-tumorigenic immortalized mammary epithelial MCF-10A cells to scrutinize its impact on cell viability and purinergic signaling.

Our results show that inhibitors of E-NTPDases, ENPPases and TNAP all significantly increased the release of eATP with chemotherapy exposure. However, only the E-NTPDase inhibitor PSB 069, a sulfonated tetracyclic compound, but not POM-1, another E-NTPDase inhibitor, consistently and significantly exacerbated chemotherapy-induced cell death. Both are inhibitors of multiple E-NTPDase isoforms. Some reports suggest that POM-1 also blocks several P2XRs. This may interfere with cell death and may explain their differing effects on chemotherapy-induced cell death. ENPPase and TNAP substrates are not limited to ATP, and therefore, inhibition of these enzymes may have ATP non-specific effects.

We also showed that the addition of exogenous eATP in the absence of chemotherapy significantly reduced TNBC cell viability. Specific inhibitors of P2RX4 and P2RX7, but not a non-specific P2RX inhibitor, attenuated the effects of ATP on cell viability and the effects of E-NTPDase inhibitors on eATP levels and their positive effects on chemotherapy-induced cell death. These data show that ATP-induced cell death and E-NTPDase-inhibitor induced augmentation of chemotherapy-induced cell death are mediated through P2RX4 and P2RX7 channels. However, two observations suggest that P2RX4 and P2RX7 activation alone may not be sufficient for the loss of cell viability observed. Firstly, the addition of eATP to cells was toxic to MDA-MB 231 and Hs 578t cells but minimally affected MDA-MB 468 cells.

However, upon chemotherapy treatment, an E-NTPDase inhibitor significantly increased eATP levels and augmented chemotherapy-induced cell death in all the TNBC cell lines. Secondly, the addition of eATP induced loss of TNBC cell viability at concentrations that were higher than those that were observed upon chemotherapy treatment in the presence of eATPase inhibitors. Thus, although P2RX4 and P2RX7 activation are necessary for the augmentation of chemotherapy-induced cell death by eATP, other factors may be required in parallel. For example, the NLRP3 inflammasome is one such death pathway whose activation is dependent on eATP but must also be primed by other factors such as NF-κB activation.

While attempting to understand the differences in concentrations required for cytotoxicity of extrinsically-applied eATP versus that of chemotherapy-induced rises in eATP, it is important to consider previous research which suggests that eATP concentrations in the immediate pericellular region may far exceed those in the bulk interstitial fluid (Pellegatti et al., 2005).

Thus, our measurements of eATP in the bulk supernatants may have underestimated pericellular eATP concentrations. In addition, ATP-induced signaling may occur in membrane demarcated intracellular organelles such as lysosomes (Qureshi, Paramasivam, Yu, & Murrell-Lagnado, 2007).

The effects of eATP on cell viability and the effects of extracellular ATPase inhibitors on eATP levels were reversed by specific P2RX4 and P2RX7 inhibitors suggesting that these receptors are not only necessary for the accentuated cell death downstream of increased eATP but also necessary for increased eATP release. The fact that P2RX4 and P2RX7 antagonists significantly attenuated eATP release even at concentrations of paclitaxel at which cytotoxicity was similar between the treatment groups, suggests that their attenuation of eATP was not due to attenuation of cell death. Given that the P2RX4 and P2RX7 antagonists but not a non-specific P2RX blocker reversed these effects, they are likely specific to these two receptor types.

We aimed to identify clinically approved compounds that modulate eATP levels. The antiparasitic drug ivermectin is an activator of P2RX4 and P2RX7 (Khakh et al., 1999; Nörenberg et al., 2012). We showed that consistent with this activity, ivermectin sensitized TNBC cell lines to chemotherapy. Interestingly, we also observed increased eATP release in chemotherapy-treated cells in the presence of ivermectin. This finding is consistent with our data indicating that P2RX4 and P2RX7 channels are not only necessary for ATP-induced loss of viability but also for eATP release.

Concerning expression levels, our western blot data show that P2RX4 is highly expressed in some TNBC cell lines as compared to immortalized mammary epithelial cells. However, our flow cytometry data revealed significantly decreased cell surface expression of P2RX4 in the TNBC cell lines as compared to between MCF-10A cells. Previous publications suggest that the majority of P2RX4 is expressed on lysosomal membranes and that cell surface expression can be increased by stimuli that induce lysosomal exocytosis such as calcium ions (Qureshi et al., 2007). This may explain the different expression patterns detected by western blot and flow cytometry.

On western blot analysis of mammary cells, we detected a specific band corresponding to the full-length glycosylated form of P2RX7 in the MDA-MB 231 and MCF-10A cells and lower molecular weight bands that may represent unglycosylated forms of P2RX7 in the MDA-MB 468 and Hs 578t cells. On the other hand, our flow cytometry data shows that P2RX7 is expressed at the cell surface of all the TNBC cell lines at higher levels than MCF-10A cells and in the presence of paclitaxel some TNBC cell lines expressed more P2RX7. This suggests there may be selection pressure for higher expression of P2RX7 in TNBC cell lines. Several published data support the facilitator role played by extracellular adenosine, derived from the metabolism of eATP, for the survival of cancer cells by inducing cell-autonomous effects on proliferation and cancer stem cell-like properties as well as paracrine effects on angiogenesis and immunoevasion (Blay, White, & Hoskin, 1997; Du et al., 2015; Fernandez-Gallardo, González-Ramírez, Sandoval, Felix, & Monjaraz, 2016; Lan et al., 2018). Given our data suggesting that P2RX7 is necessary for eATP release, this may explain selection for higher levels of expression. Although expression levels may be low, given that a specific inhibitor of P2RX7 markedly attenuated the cytotoxic effects of eATP and attenuated the positive effects of E-NTPDase inhibitors on eATP levels and loss of cell viability upon chemotherapy exposure, it is possible that even low levels of expression P2RX7 may have functional consequences due to the formation of non-selective macropores in the cell membrane.

## Author Contributions

All authors contributed to review and analysis. JM performed a majority of the assays with NW executing the RNA analysis and JD carrying out the Western blot analysis. JM and MC conceived of and designed the experiments, reviewed the data, authored and edited the manuscript.

