## Supplemental Data for "Augmentation of extracellular ATP synergizes with chemotherapy in triple negative breast cancer"

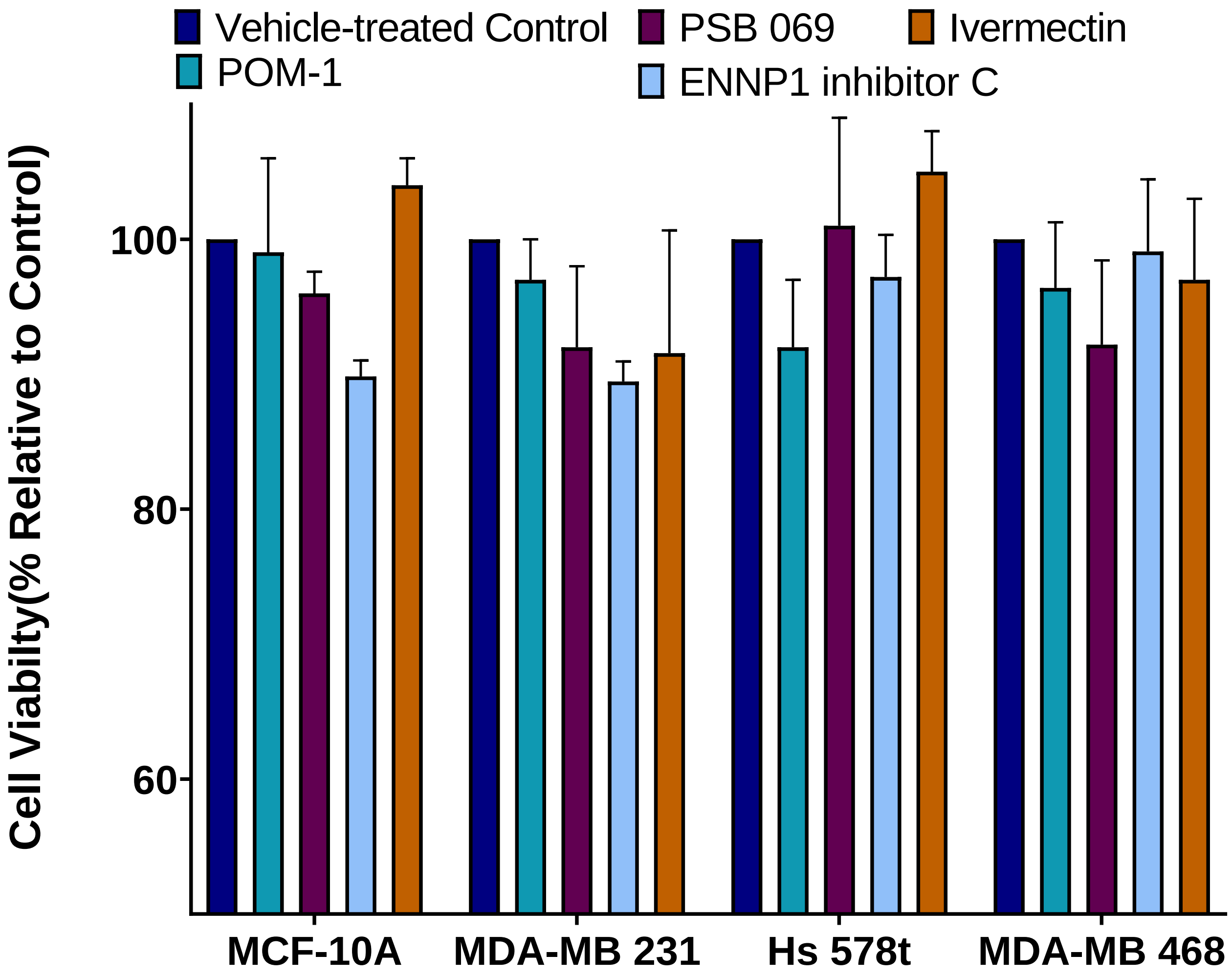


**Figure S1: Cell viability of eATPase inhibitors**. TNBC and MCF-10A cell lines were treated with POM-1 (E-NTPDase inhibitor, 10 µmol/L), PSB 069 (E-NTPDase inhibitor, 10 µmol/L), ENNP1 inhibitor C (ENPP1 inhibitor, 10 µmol/L) or vehicle addition for six hours, and cell viability was measured using the PrestoBlue HS assay. Standard deviation was calculated from three independent experiments performed in triplicate. Y axis set at 50%.

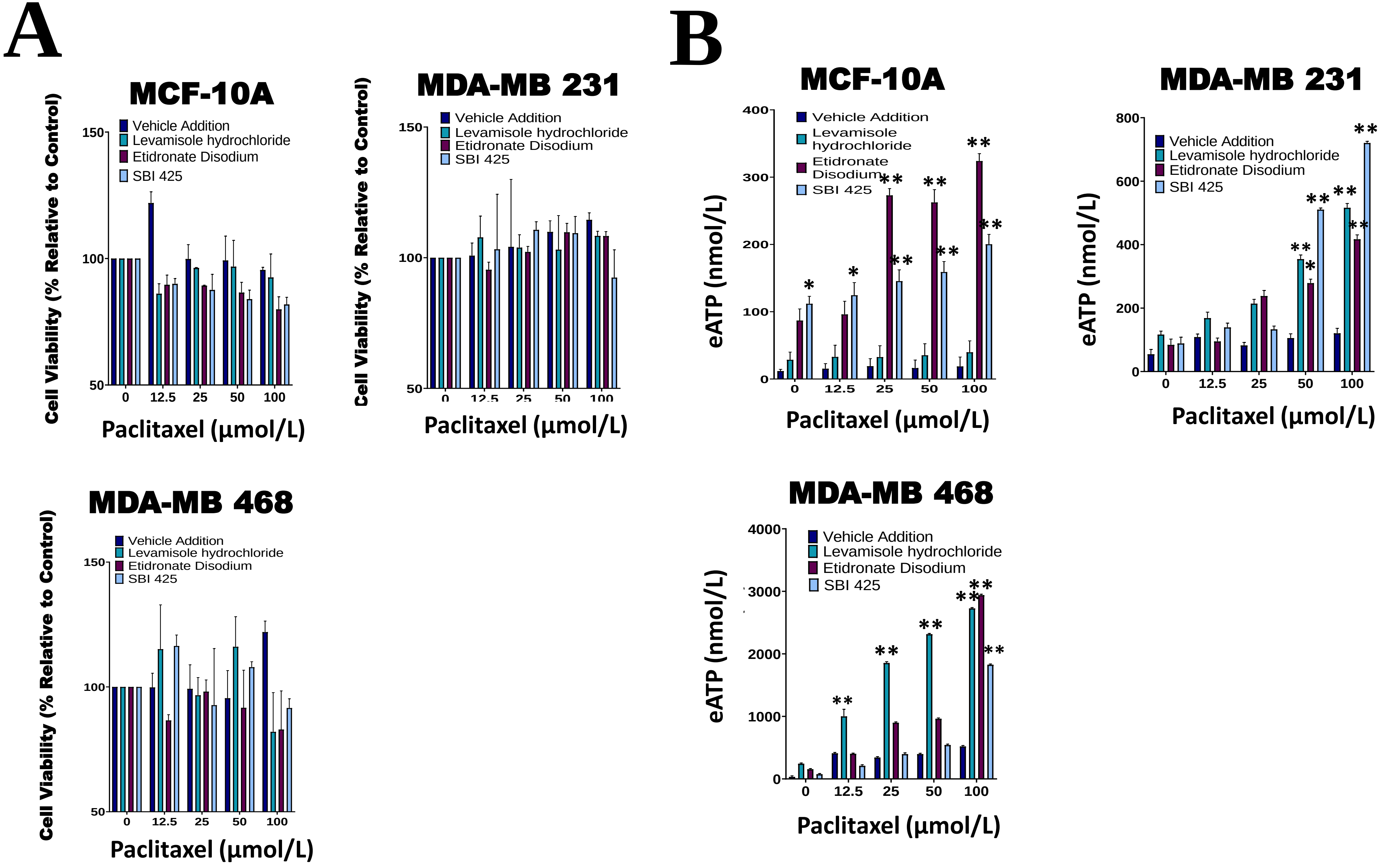


**Figure S2: Comparing eATP release from paclitaxel-treated cells in the presence of phosphatase inhibitors or vehicle addition. (A)** TNBC cell lines and MCF-10A cells were treated for six hours with phosphatase inhibitors levamisole hydrochloride (tissue non-specific alkaline phosphatase inhibitor, 50 µmol/L), SBI 425 (tissue non-specific alkaline phosphatase inhibitor, 10 µmol/L), etidronate disodium (protein tyrosine phosphatase inhibitor, 50 µmol/L) or vehicle addition, and cell viability was measured with PrestoBlue HS. Standard deviation was calculated from three independent experiments performed in triplicate. (**B)** eATP concentrations were measured in the supernatants of paclitaxel-treated TNBC cell lines and MCF-10A cells in the presence of the phosphatase inhibitors levamisole, SBI 425, etidronate sodium, or vehicle addition with paclitaxel. Y axis set at 50%. Student's t-test was performed to ascertain significance. * represents $p<0.05$ and ** represents $p<0.01$.

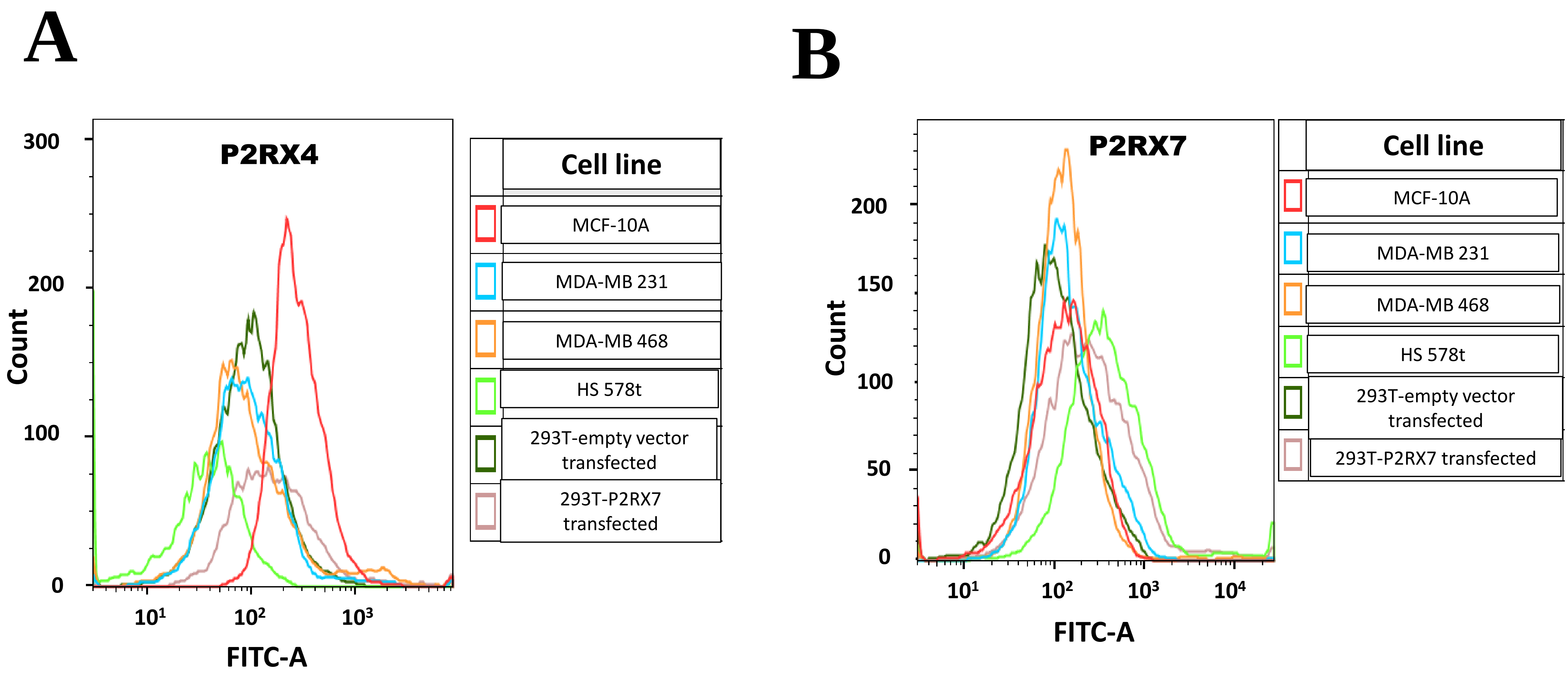


**Figure S3: Cell surface expression analysis of P2RX4 and P2RX7.** Histograms displaying the different extracellular (**A)** P2RX4 and (**B)** P2RX7 expressions in TNBC cell lines, MCF-10A cells, and HEK 293T cells transfected with P2RX4 or P2RX7 as positive controls.

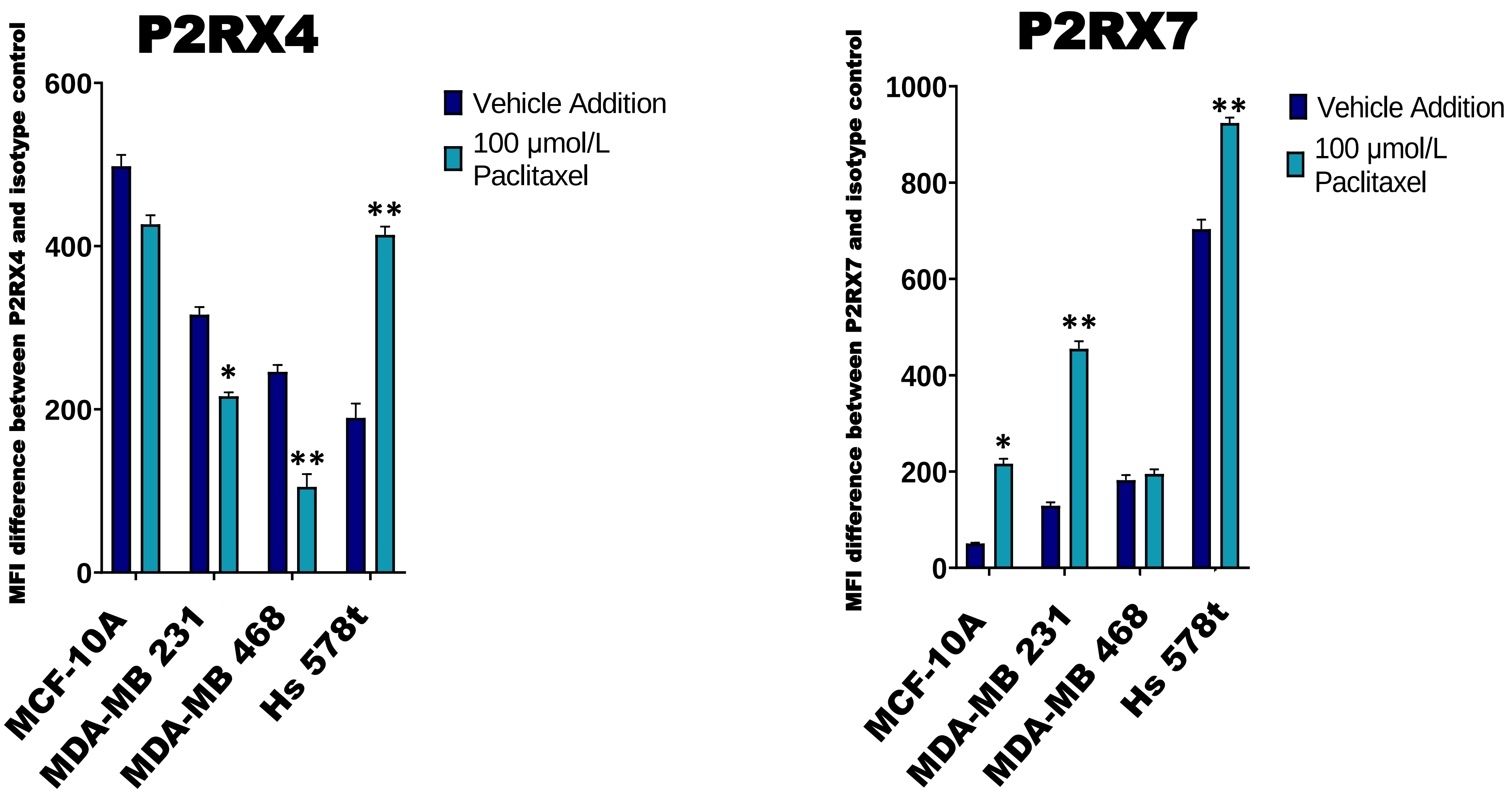


**Figure S4: Paclitaxel's impact on the cell surface expression of P2RX4 and P2RX7.** The calculated difference in mean fluorescence intensity (MFI) values between TNBC cell lines and MCF-10A cells in the absence of presence of 100 µmol/L paclitaxel stained with **(A)** P2RX4 or **(B)** P2RX7 specific antibody and the isotype control for the different cell lines examined. Student's t-test was performed to ascertain significance. * represents $p<0.05$ and ** represents $p<0.01$ relative to MFI difference in MCF-10A cells.
